# Membrane positioning across antigen-induced synaptic contacts tunes CAR-T cell signaling and effector responses

**DOI:** 10.1101/2023.10.01.560371

**Authors:** Fenglei Li, Kaushik Choudhuri

## Abstract

Tumor antigen recognition by chimeric antigen receptors (CAR) triggers phosphorylation of their cytoplasmic portions resulting in CAR-T cell activation. We and others have shown that immunoreceptor triggering depends on the formation of close synaptic contacts, determined by the span of immunoreceptor-ligand complexes, from which large inhibitory phosphatases such as CD45 are sterically excluded. Here, we show, varying CAR-antigen complex span, that CAR-T cell activation depends on a formation of close contacts with target cells. CAR-antigen complexes with a span of 4 immunoglobulin superfamily (IgSF) domains maximize CAR-T cell activation, closely matching the span of endogenous TCR-pMHC complexes. Longer CAR-antigen complexes precipitously reduced triggering and cytokine production, but notably, anti-tumor cytotoxicity was largely preserved due to a ∼10-fold lower signaling threshold for mobilization of cytolytic effector function. Increased intermembrane spacing disrupted short-spanned PD-1-PD- L1 interactions, reducing CAR-T cell exhaustion. Together, our results show that membrane positioning across the immunological synapse can be engineered to generate CAR-T cells with clinically desirable functional profiles *in vitro* and *in vivo*.

## Introduction

T cells engineered to express synthetic chimeric antigen receptors (CAR) are a promising new modality for cancer immunotherapy (Boyiadzis et al., 2018). Despite prominent successes in treating B cell ‘liquid tumor’ malignancies (Maude et al., 2014; Porter et al., 2011; Schuster et al., 2017), significant challenges to more widespread adoption remain, especially against solid tumors. CAR-T cell immunotherapy is limited by frequent treatment-related adverse effects, including cytokine release syndrome, which can be life-threatening, and treatment failures due to CAR-T cell exhaustion, or trogocytic tumor antigen extraction (Morris et al., 2022).

Second generation CARs currently in clinical use target tumor antigens *via* an extracellular scFv antibody fragment, tethered to the T cell surface by structurally heterogeneous spacers (June et al., 2018). CAR intracellular portions consist of a membrane-proximal costimulatory domain, derived from CD28 or 4-1BB, linked to immunoreceptor tyrosine-based activation motifs (ITAMs) derived from the T cell antigen receptor (TCR) ζ chain. Tumor antigen binding results in CAR triggering *via* phosphorylation of its cytoplasmic ITAMs. It has long been known that CAR-T cell activation is exquisitely sensitive to the size of CAR spacers (Guedan et al., 2019; Hudecek et al., 2013; Hudecek et al., 2015; Stoiber et al., 2019). We and others have shown that triggering by native TCRs and chimeric immunoreceptors occurs through binding-induced segregation of large cell-surface phosphatases, such as CD45, from engaged receptors at T cell synaptic contacts, allowing stable phosphorylation of engaged immunoreceptors at synaptic close contacts zones (Choudhuri et al., 2009; Choudhuri et al., 2005; Cordoba et al., 2013; Xiao et al., 2022). CAR triggering is abrogated by increasing the CAR-antigen complex span, by insertion of rigid membrane-proximal spacers into the CAR ectodomain, which results in increased intermembrane separation at CAR-T cell synaptic contacts with tumor cells, and greater access of CD45 to engaged immunoreceptors – in keeping with the kinetic-segregation model of immunoreceptor triggering.

Here, we extend these findings to CAR triggering by showing that increasing CAR-antigen dimensions increases intermembrane spacing at CAR-T cell synapses, enabling greater CD45 access to engaged CARs. While this severely attenuates cytokine secretion, anti-tumor cell cytotoxicity is largely preserved due to ∼10-fold lower signaling threshold for mobilizing T cell lytic machinery, relative other effector functions.

## Results

### Increases in CAR-antigen span attenuates triggering by predictably increasing intermembrane separation and CD45 access at CAR-T cell synapses

To test whether CD45 segregation form CARs at synaptic contacts affected CAR triggering according we inserted xenogeneic immunoglobulin superfamily (IgSF) containing spacers, derived from the complete ectodomains of rat Thy-1 and mouse CD4, into the membrane-proximal stalk of a GFP-targeting CAR (αGFP-0CAR or 0D CAR). The 0D CAR comprised of a camelid VHH domain specific for GFP tethered to the plasma membrane by a short flexible stalk, transmembrane segment and cytoplasmic signaling domains of human CD28, fused to the intracellular domain of the T cell antigen receptor ζ chain (TCRζ) (Fig. 1A). A CAR with elongated ectodomain was generated by insertion of a spacer, derived from the complete ectodomain of mouse CD4, into the membrane proximal region of the 0D CAR ectodomain (αGFP-CD4 CAR). CAR constructs were expressed in purified polyclonal CD8+ T cells, isolated from human donors, by lentiviral transduction. We also generated, by lentiviral transduction, CHO cells stably expressing a membrane-tethered version of GFP with the rat Thy-1 ectodomain inserted into the membrane-proximal stalk region to serve as a target cell (GFP-Thy1). Cells were sorted for comparable expression by flow-cytometry to directly compare responses between CAR-T cells expressing 0D and αGFP-CD4 CARs (Fig 1B).

**Figure 1.**
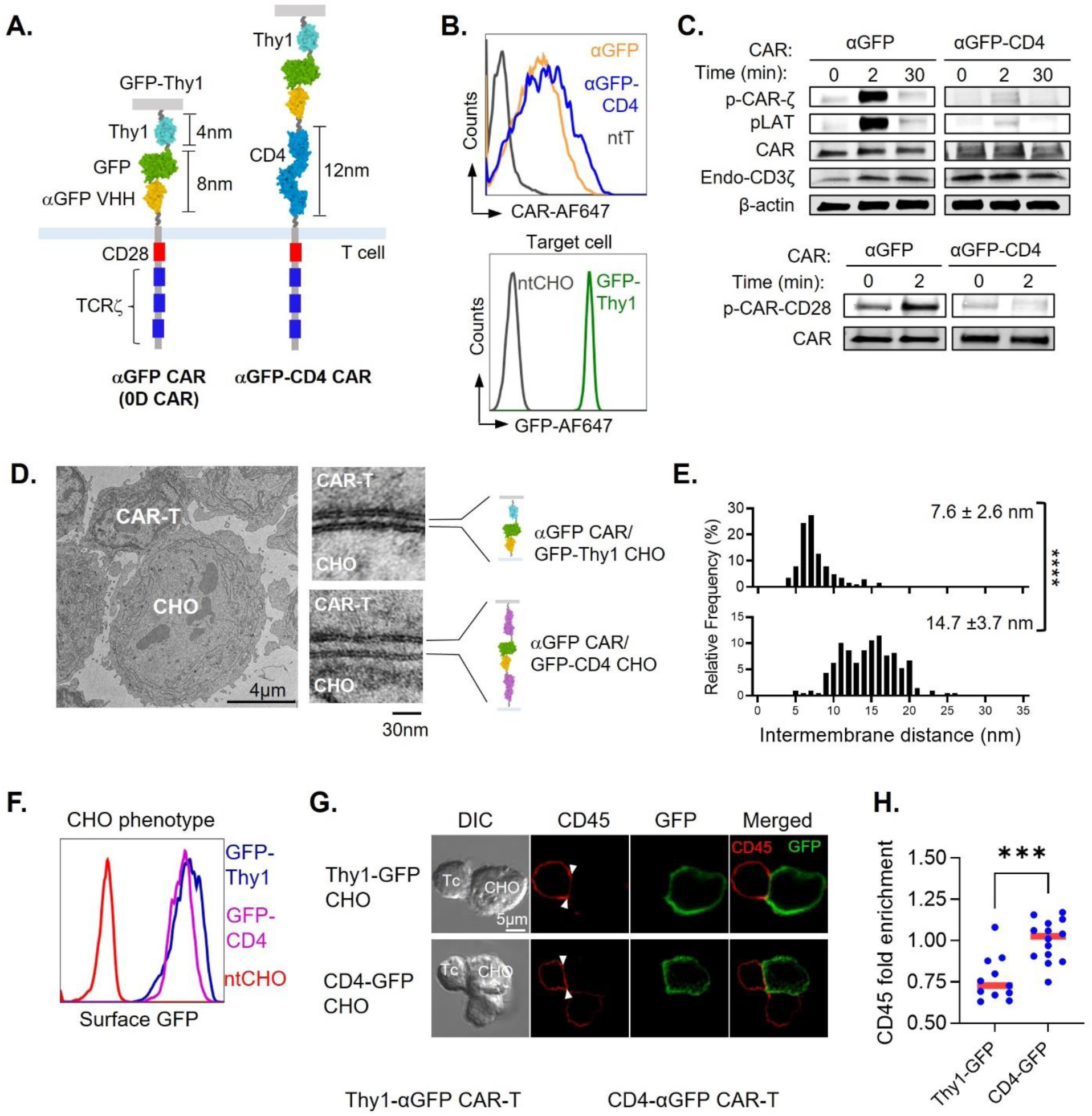
CAR ectodomain elongation attenuates proximal signaling by increasing intermembrane distance and CD45 access at synaptic contacts. (A) Schematic of a GFP-targeting 2^nd^ generation CAR construct (αGFP/0D CAR), and an elongated version in which the full ectodomain of mouse CD4 is inserted membrane-proximally to the CAR VHH antibody domain (αGFP-CD4 CAR). Both CARs have identical cytoplasmic portions containing the human CD28 signaling domain followed by three ITAMS of human TCRζ. CARs were expressed in human polyclonal CD8+ T cells (CAR-T cells), and are shown in complex with model target antigen GFP-Thy1 expressed in CHO cells. GFP-Thy1 consist of GFP linked to a rat Thy-1 membrane-proximal spacer, tethered to the plasma membrane by a short CD28 (aa561- 609) spacer and transmembrane segment. Molecular complexes are shown approximately to scale, with indicated dimensions measured from crystallographic structures. (B) Histogram of surface expression levels of the indicated CARs in CAR-T cells sorted for comparable expression levels (*top panel*), and surface levels of GFP-Thy-1 expressed on CHO target cells (*bottom panel*). (C) *Top panel*, Immunoblot of CAR-associated TCRζ ITAM phosphorylation (pCAR-ζ) and tyrosine phosphorylation of linker for activation of T cells (pLAT^Y191^) in CAR-T cells expressing the indicated CARs. Samples were briefly centrifuged to initiate contact and kept on ice prior to incubation at 1:1 ratio with CHO cells expressing GFP-Thy1 for 2 and 30 minutes at 37°C. Samples were lysed on ice for the 0 minute timepoint. Immunoblots of total CAR-associated TCR-ζ and β-actin are shown as expression and loading controls. Results are representative of 4 independent experiments. *Bottom panel*, Immunoblot of tyrosine phosphorylation of CAR-associated CD28 signaling domain in CAR-T cells expressing the indicated CARs and treated as in *Top panel*. Results are representative of 3 independent experiments. (D) *Left panel*, Representative transmission electron micrograph of CAR-T cell (expressing αGFP CAR) conjugated with a GFP-Thy-1-expressing CHO cell. *Right panels*, Representative high magnification images of apposed plasma membranes at synaptic contacts between CAR-T cells expressing the indicated CARs and CHO cell expressing either GFP-Thy1 or GFP-CD4. (E) Normalized frequency histograms of measurements of intermembrane distance along the synaptic interface between indicated CAR-T cells and CHO cells expressing GFP-Thy1 or GFP-CD4. n=205 measurements at αGFP CAR/GFP-Thy1 CHO interfaces from 17 conjugates, and 209 measurements at αGFP CAR/GFP-CD4 CHO interfaces from 23 conjugates, pooled from 3 independent experiments. Means ± s.d are shown. Means were compared using two-tailed *t*-test. ****, P < 0.0001. (F) Histogram overlay of surface expression levels of CHO cells expressing GFP-Thy1 or GFP-CD4 sorted for comparable expression levels. Untransfected CHO cells (ntCHO) are shown as a labeling control. (G) Representative differential interference contrast (DIC) and confocal fluorescence images of conjugates of 0D-CAR-T cells with CHO cells expressing GFP-Thy1 or GFP-CD4. Arrowheads indicate the synaptic contact interface. Cells fixed, and permeabilized prior to staining with antibodies against CD45 (red) and GFP (green), and appropriate fluorescently labeled secondary antibodies. (H) Quantitation of CD45 and GFP colocalization at contact interfaces by Pearson’s correlation coefficient (PCC). Data points represent measurements from individual interfaces. Means were compared using two-tailed *t*-test. ****, P < 0.005.

We assessed triggering by 0D and elongated CARs by immunoblot of phosphorylated TCRζ and downstream signaling adapter linker for activation of T cells (LAT). CAR-T cells and CHO were gently centrifuged to initiate contact, and either lysed on ice, or incubated for 2 or 30 minutes at 37°C. 0D CAR was robustly phosphorylated at 2 min incubation, which returned to near-baseline levels by 30 minutes (Fig 2C). In contrast, αGFP-CD4 CAR-T cells showed minimal increase in CAR phosphorylation, barely above background levels.

**Figure 2.**
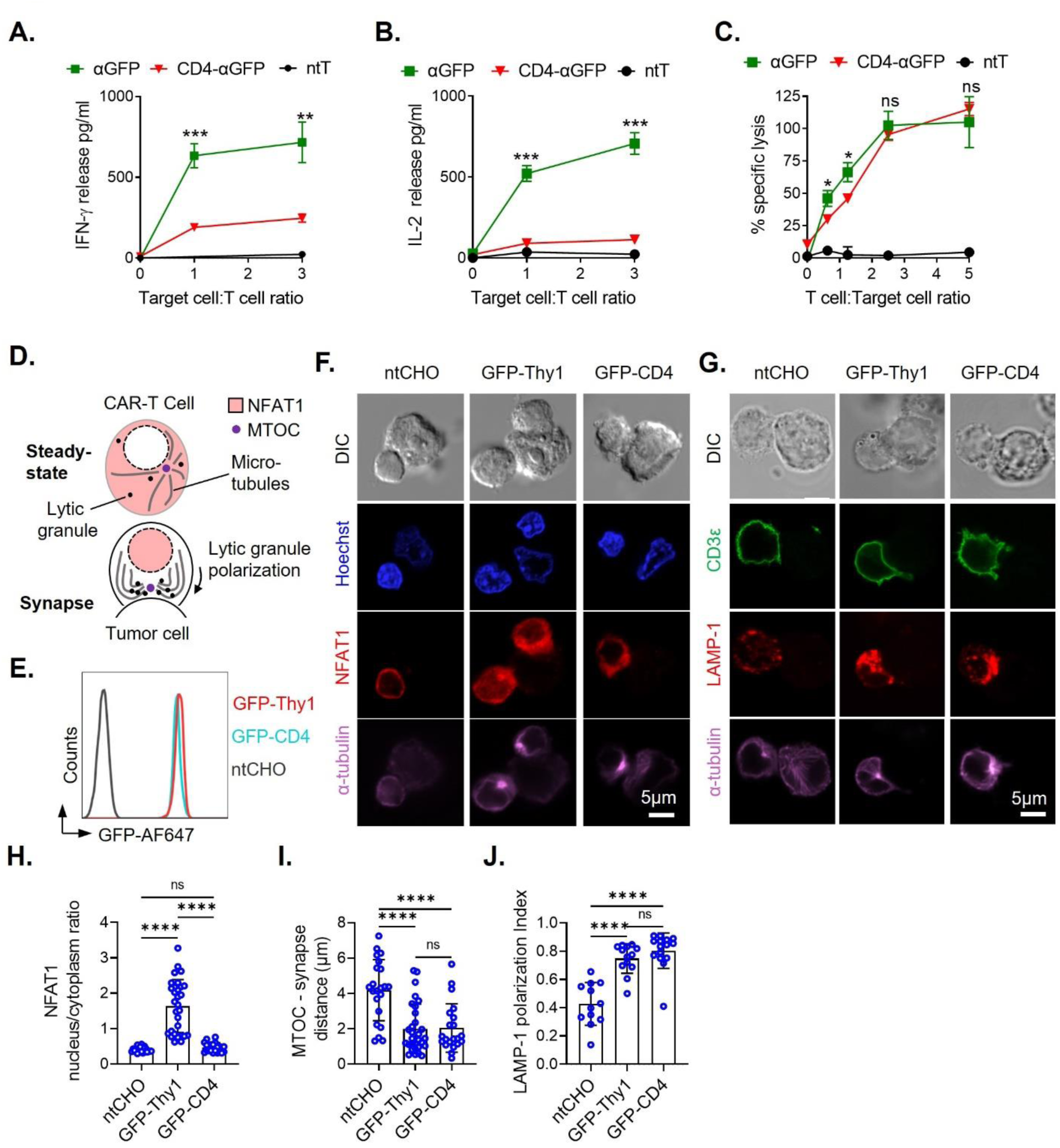
Hierarchical cell-intrinsic signaling thresholds differentially engage CAR-T cell cytotoxicity and transcriptional activation. (A) IFNγ and (B) IL-2 release by indicated CAR-T cells (10^4^/well) cocultured for 24hrs with increasing numbers of GFP-Thy1-expressing CHO target cells, shown as Target cell/T cell ratio. The indicated cytokines (IFNγ and IL-2) were measured by ELISA of culture supernatants. Mean ± s.d are shown for each data point. Results are representative of 4 independent experiments. (C) CAR-T cell-mediated cytotoxicity in cocultures of the indicated CAR-T cells with BATDA-loaded CHO cell targets expressing Thy1-GFP. BATDA released to the culture supernatants, as a measure of CHO cell lysis, was detected using fluorescent chelates formed in europium solution and time-resolved fluorescence. Data are pooled from two independent experiments. Data points are mean ± s.d. (D) Schematic depicting CAR-T cell microtubule organizing center (MTOC) and lytic granule polarization to synaptic contacts with tumor cell targets. (E) Histogram overlay of surface expression levels of GFP-Thy1 and elongated GFP-CD4 on CHO cells sorted for comparable expression levels. ntCHO, untransduced CHO cells. (F) Representative confocal fluorescence images of conjugates between CAR-T cells, expressing αGFP CAR, and CHO cells expressing GFP-Thy1 or GFP-CD4 target antigens. Shown are differential interference contrast (DIC) images of CAR-T :CHO cell conjugates, and corresponding fluorescence channels of DAPI-labeled dsDNA, and specific antibody-labeled NFAT1c transcription factor and α-tubulin in microtubules, both detected using appropriate fluorescently-tagged secondary antibodies. (G) Representative confocal images of CAR-T cell conjugates as in *(F)* labeled with specific antibodies and appropriate fluorescently-tagged secondary antibodies against CAR-T cell CD3ε, lytic granule marker LAMP-1 and α-tubulin in microtubules. (H) Quantitation of NFAT1 activation as measured by cytoplasm-to-nucleus translocation using confocal microscopy as in *(F)*. Results are expressed as a ratio of mean NFAT fluorescence intensity in CAR-T cell nucleus/cytoplasm. Bars are means ± s.d., and data points represent individual cells. Results are representative of 4 independent experiments. (I) Quantitation of MTOC recruitment to the CAR-T cell synapse as measured by distance (μm) of the MTOC (as imaged in *(F)* and *(G)*) from the CAR-T cell:Target cell contact interface. Data points represent measurements from individual cells. Results are representative of 4 independent experiments. Bars are mean ± s.d., and data points represent individual cells. Results are representative of 4 independent experiments. (J) Quantitation of cytotoxic granule polarization towards the CAR-T cell synapse (measured using the granule marker LAMP-1 as in *(G)*. Bars are mean ± s.d., and data points represent individual cells. Data are pooled from 2 independent experiments. Means were compared by one-way ANOVA corrected for multiple comparisons. ****, P < 0.0001; ***, P < 0.001; **, P < 0.01; ns, not significant (P > 0.05).

We next measured intermembrane distances at contact interfaces between GFP-Thy1-expressing CHO cells, and CAR-T cells by transmission electron microscopy (TEM). We measured distances between apposed membranes along the entire interface in regions where the membranes were parallel and exhibited a tram-line appearance, indicating that they were positioned orthogonally to the sectioning plane (Fig. 1D). For these experiments we employed 0D CARs conjugated to CHO cells expressing GFP-Thy1, or an elongated version with the mouse CD4 ectodomain in the membrane-proximal stalk (GFP-CD4). The mean intermembrane distance at contact interfaces of conjugates between GFP-Thy1 CHO and 0D CAR-T cells was ∼ 8 nm, while that at interfaces with CD4 CAR was significantly increased to ∼15 nm, demonstrating that elongating the CRA ectodomain increases intermembrane separation at synaptic contacts (Fig. 1E).

To investigate the distribution of CD45 at synaptic contacts we made conjugates between CHO cells and CAR T cells and labeled CD45 and on T cells and GFP (as a proxy for engaged CARs) at contact interfaces of CAR-T cell-CHO cell conjugates by confocal microscopy (Fig. 1F,G). We analyzed contact interfaces for colocalization between CD45 and GFP by Pearson’s correlation coefficient (PCC), which demonstrated low PCC (indicating relative segregation/dispersion) between CD45 and GFP, and therefore 0D CAR by proxy, while conjugates with CHO cells expressing GFP-CD4 showed a highly significant increase in PCC, indicating greater colocalization of GFP-CD4 with CD45 at interfaces, indicating abrogation of CD45 segregation (Fig. 1H).

### Hierarchical signaling thresholds elicit CAR-T cell cytotoxicity and transcriptional activation

We next tested whether reduction in triggering due to CAR elongation affected CAR-T cell activation, by measuring cytokine secretion and target cell killing. We co-cultured 10^4^ 0D or CD4 CAR-T cells with increasing numbers of target cells (presented as a Target:T cell ratio) expressing GFP-Thy1 for 24 hours, and measured IFNγ and IL-2 release in culture supernatants by ELISA.

To assess CAR-T cell mediated killing we utilized the DELFIA time-resolved fluorescence cytotoxicity assay, in which target cells were loaded with lanthanide-chelator BATDA that fluoresces when bound to lanthanides such as Europium. Loaded target cells were incubated with increasing numbers of CAR-T cells (presented as a T cell: Target cell ratio) for 2 hrs, BTADA release due to cell killing and quantitated by measuring its fluorescence following addition of Europium to samples, by time-delayed fluorimetry (Blomberg et al., 1996). 0D CAR T cells robustly responded to GFP-Thy1 target cells by releases IFNγ and IL-2 in a titratable manner (Fig. 2A,B), which approached saturation at high T cell: Target cell ratio. In stark contrast, cytokine secretion by CD4 CAR-T cells was severely blunted, reaching ∼25-30% of 0D CAR-T cell levels at the highest Target:T cell ratio (Fig. 2A,B). Surprisingly, CAR-T cell-mediated cytotoxicity against target cells was largely preserved by CAR elongation, with a reduction in cell killing only apparent, and significantly different at low T cell: Target cell ratios (Fig. 2C).

To investigate the cell biological basis for these differences in cytokine release and cytotoxic effector responses, we measured microtubule organizing center (MTOC) and lytic granule recruitment (Fig. 2D, E)of 0D CAR-T cells to the synaptic contact with GFP-Thy1 and GFP-CD4 expressing target cells (Fig. 2G), to detect activation of lytic machinery (Jenkins et al., 2009), and nucleocytoplasmic shuttling of the transcription factor NFAT1, as a marker of transcriptional activation (Fig. 2D, F) (Neilson et al., 2001; Patra et al., 2004). In keeping with functional response, NFAT1 shuttling to the CAR-T cell nucleus was severely affected, remaining at baseline levels (in conjugates with untransfected CHO cells – ntCHO) (Fig. 2H), while MTOC polarization (Fig. 2I), quantitated as MTOC distance from synapse, and secretory granule polarization (Fig. J), were unaffected by CAR-antigen complex elongation. Taken together, these results demonstrate a hierarchy in signaling thresholds for CAR-mediated effector responses, in which T cell cytotoxicity is triggered at much lower levels of CAR signaling than that required for transcriptional activation and cytokine production.

### Increased intermembrane separation disrupts PD-1-PDL1 interactions

We reasoned that increasing intermembrane separation at CAR-T cell synaptic contacts, by elongating CAR-antigen complexes, might affect other key receptor-ligand interactions with short spans, that signal at the CAR-T cell immunological synapse. An important negative regulator of synaptic signaling is the small, 1 immunoglobulin superfamily domain (IgSF)-containing receptor PD-1, which binds its 2IgSF domain-containing ligand PD-L1 on target cells (Zak et al., 2017). Ince PD-1 is only transiently upregulated in activated T cells, we chose to overexpress PD-1 in CAR-T cells by lentiviral transduction ((Fig. 3A). To asses synapse formation and PD-L1 engagement of CAR-T cells we first monitored 0D PD-1+ CAR-T cells forming synapses on supported lipid bilayers (SLB) containing GFP-His10, and fluorescently-labeled PD-L1 and ICAM-1. CAR and PD-1 distributions at synapses was inferred by enrichment and clustering of GFP and PD-L1 on bilayers.

**Figure 3.**
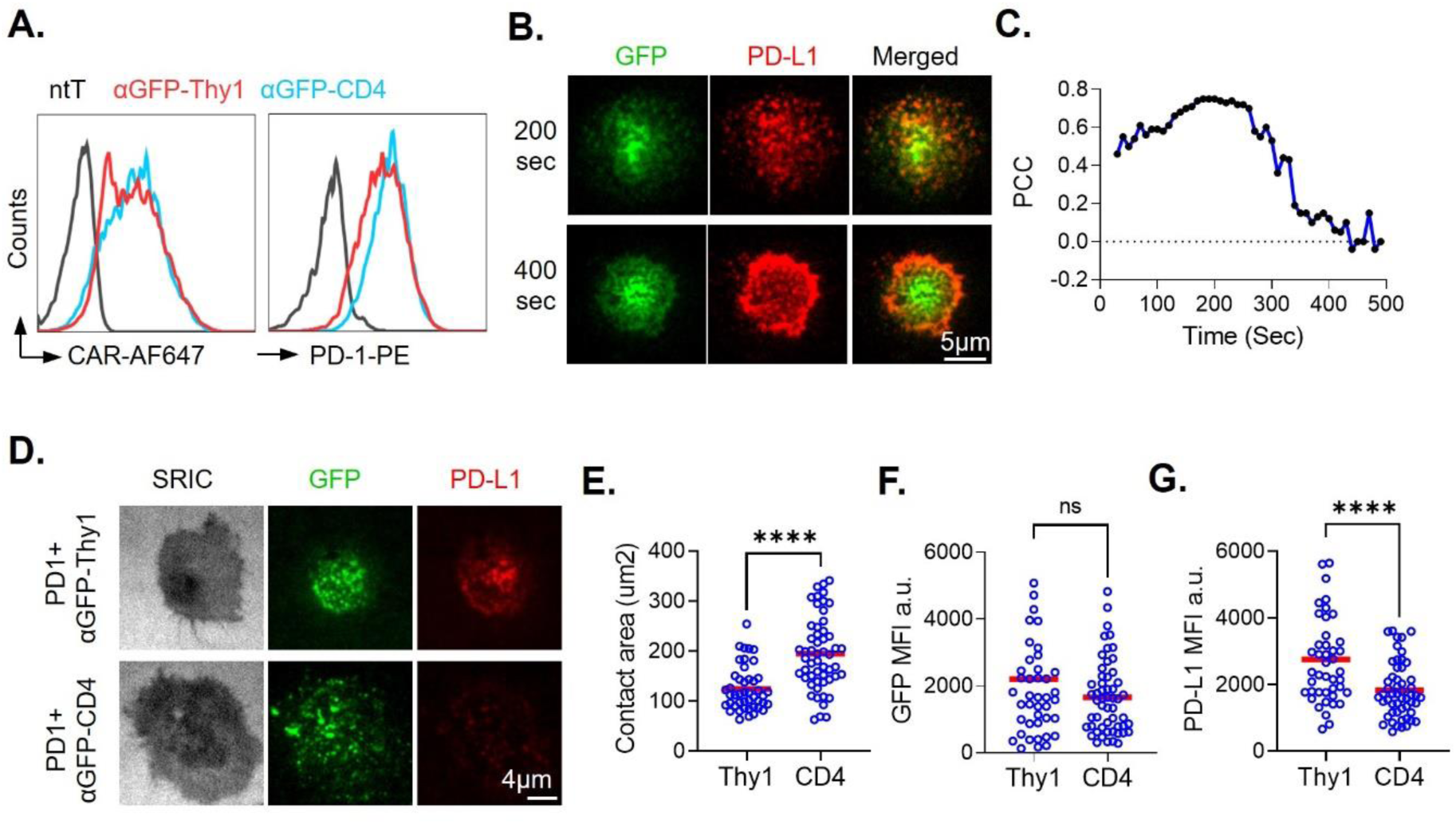
Increased CAR-antigen complex span disrupts PD-1 engagement. (A) Histogram overlays of surface levels of the indicated CARs (*left panel*) and PD-1 (*right panel*) expressed in CAR-T cells, sorted for comparable expression and measured by flow-cytometry. (B) Representative TIRFM images of GFP and PD-L1-AF647fluorescence of human CD8+ T cells transduced with αGFP-Thy1 CAR and PD-1 interacting with SLB containing 200 molecules/μm^2^ of GFP, 300 molecules/μm^2^ of ICAM-1, and 100 molecules/μm^2^ of PD-L1-AF647 Images are shown of the same CAR-T cell synapse formed on SLB at 200s after initial contact (top panels) and 400s after contact with bilayers. (C) Pearson’s correlation coefficient (PCC) was calculated as a measure of colocalization between GFP fluorescence, as a proxy for engaged αGFP-Thy1 CAR distribution, and fluorescently labeled PD-L1, as a proxy for PD-1 distribution at synapses. The cell (as shown in *B*.) were monitored at 10s intervals and each data point indicates the PCC at the cell synapse for the indicated timepoint. Dotted line indicates 0 PCC (uncorrelated distribution). Results are representative of 4 independent experiments. (D) Representative TIRFM images of GFP and PD-L1-AF647 fluorescence of CAR-T cells transduced with the indicated CARs and PD-1, interacting with SLB constituted as in *(B)*. SRIC, surface reflection interference contrast image. (E) Quantitation of synapse contact area 120s after initial contact with SLBs (measured form SRIC images) of CAR-T cells expressing αGFP-Thy1 (Thy1) or αGFP-CD4 (CD4) and PD1. Results are representative of 4 independent experiments. (F) and (G) Quantitation of GFP (F) and PD-L1-AF647 (G) fluorescence intensity at synapses as in *(D)*. MFI, mean fluorescence intensity; a.u. arbitrary units. Results are representative of 3 independent experiments. Red bars represent means and data points represent measurements from individual cells. Means were compared using two-tailed *t*-test. ****, P < 0.0001; ns, not significant (P > 0.05).

As observed in native T cells interacting with cognate pMHC ligands (Varma et al., 2006), CARs formed microclusters within seconds of contacts with SLB, which were transported medially, and accumulated at the synapse center over time. PD-1 also formed microclusters, which briefly partially colocalized with CAR microclusters, but segregated from CARs within minutes, and coalesced in a peripheral domain with no overlap with CARs accumulated at the synapse center (Fig. 3B), quantitated as PCC over time (Fig. 3C). We next investigated whether CAR elongation affected binding-induced PD-1 accumulation at the synapse. In contrast to 0D CAR-T cells, CD4 CAR-T cells failed to bind and enrich PD-L1, resulting in barely detectable enrichment of PD-L1 at synapses (Fig. 3D. G). This was not due to poor adhesion or GFP engagement as synaptic contact areas were significantly larger for CD4 CAR-T cells (Fig. 3E), while they accumulated GFP comparably to 0D CAR T cells (Fig 3F).

### CARs elongated with titin-derived spacers reduce acute cytokine release *in vivo*

To test whether the observed effects of CAR elongation also occur *in vivo*, we established a leukemia/lymphoma model in NSG mice, by infusing Raji B cells, stably expressing a membrane-tethered version of GFP-Thy1 (as in CHO cells) and luciferase to monitor tumor burden by luciferin administration and intravital bioluminescence imaging (IVIS). We also generated a new elongated CAR by insertion of 3 contiguous IgSF domains of the sarcoplasmic protein titin (I64- 66) into the membrane-proximal stalk of 0D CAR (3D CAR) (Fig. 4A). This was done to preclude the possibility of any confounding interactions in *trans* between the mouse CD4 spacer and NSG mouse. We noticed in our triggering experiments that, in addition to reduction in CAR phosphorylation, the CD28 costimulatory domain also failed to be phosphorylated. While this had minimal effects on CAR proximal signaling or effector responses up to 24hr of measurement in our signaling and cytokine release assays, lack of adequate costimulation could affect CAR-T cell survival, that would complicate interpretation of the effects of CAR elongation. We therefore replaced the CD28 costimulatory domain with that of 4-1BB, which provides strong costimulation to CAR-T-cells, but does not depend on phosphorylation for signaling (Fig. 4A).

**Figure 4.**
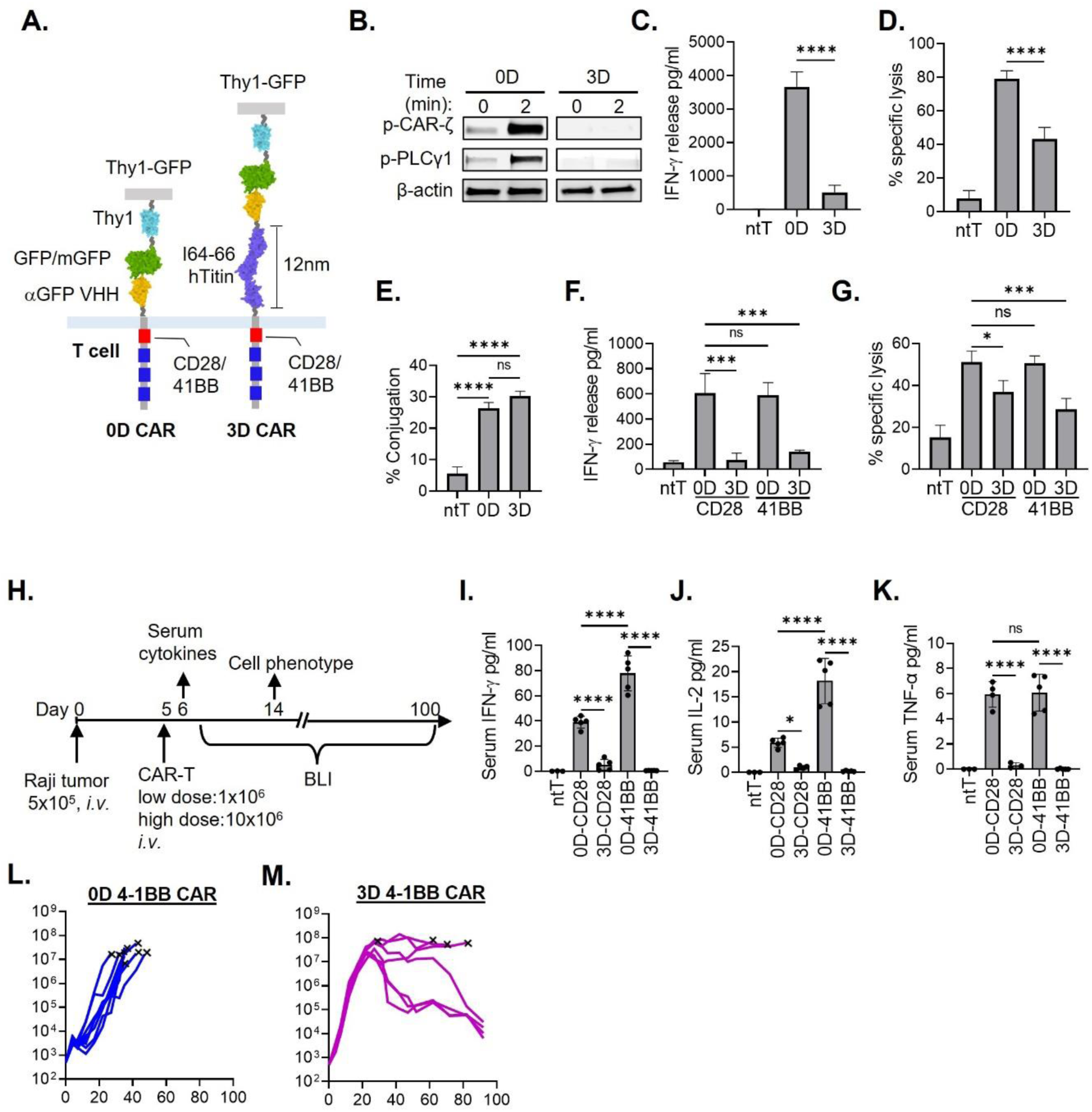
Increased CAR-antigen complex span improves CAR-T cell functional phenotype and tumor control, resulting in prolonged survival in a B cell leukemia/lymphoma model. (A) Schematic of αGFP CARs with ectodomains containing no spacer (0D) or 3 IgSF titin (3D) spacers, and cytoplasmic costimulatory domains derived from either CD28 or 4-1BB. CARs are shown in complex with Thy1-GFP target antigen expressed on CHO cells. Titin dimensions are indicated, as measured from crystallographic structures (PDB entry number: 3B43). (B) Immunoblot of tyrosine-phosphorylated CAR ITAMs (p-CAR-ζ) and phospholipase Cγ1 (p-PLCγ1) in lysates of the CAR-T cells expressing the indicated CARs incubated with CHO cells expressing Thy-1-GFP. Cells were mixed at 1:1 ratio and centrifuged briefly in the cold to initiate contact. Cells were then placed on ice, and either lysed, or incubated for 2 minutes at 37°C. β-actin is shown as a loading control. Results are representative of 5 independent experiments. (C) IFNγ release in culture supernatants, as measured by ELISA, of CAR-T cells expressing the indicated CARS cocultured with CHO cells expressing Thy1-GFP for 24hrs. Mean ± s.d. are shown. Results are representative of 3 independent experiments. (D) Specific lysis of Thy1-GFP-expressing CHO cells after 2 hrs 1:2 coculture with the indicated CAR-T cells. Mean ± s.d. are shown. Results are representative of 3 independent experiments. (E) Adhesion of the indicated CAR-T cells with Thy1-GFP-expressing CHO cells, following centrifugation and coculture at 37°C for 10 min. CAR-T cells were loaded with red fluorescent dye, CHO cells had green fluorescence as a result of the GFP expression, and percentage ‘double-positive’ cells as measured by flow cytometry taken as a measure of cell-cell adhesion. Mean ± s.d. are shown. Results are representative of 3 independent experiments. (F) Comparison of IFNγ release in culture supernatants, as measured by ELISA, of CAR-T cells expressing the 0D and 3D CARS, with either CD28 or 4-1BB costimulatory domains, cocultured for 24hrs with CHO cells expressing Thy1-GFP. Mean ± s.d. are shown. Results are representative of 5 independent experiments. (G) Comparison of specific lysis of Thy1-GFP-expressing CHO cells by CAR-T cells expressing the 0D and 3D CARS, with either CD28 or 4-1BB costimulatory domains, cocultured for 2hrs with CHO cells expressing Thy1-GFP. Mean ± s.d. are shown. Results are representative of 5 independent experiments. (H) Schematic of human Raji B cell leukemia/lymphoma tumor model in NOD-*scid* IL2Rgamma^null^ (NSG) mice. Mice were seeded *i.v.* with the indicated number of firefly luciferase-expressing Raji B cells at Day 0, and indicated numbers of CAR T cells (low-and high-dose) introduced *i.v.* on Day 5. Serum was collected 24hrs after high dose CAR-T cell infusion (Day 6) to measure cytokine levels. In separate experiments, mice were infused with low-dose CAR-T cells and followed for survival and tumor burden monitored by bioluminescence imaging (BLI) every 5 days after. Cells were collected for phenotyping by flow-cytometry at Day 14. (I)-(K) Measurement of serum cytokines IFNγ (I), IL-2 (J) and TNFα (K), 24hrs following high-dose infusion of the CAR-T cells expressing the indicated CARs. Each data point represents one mouse. Means ± s.d are shown. Results are representative of 3 independent experiments. (L, M) BLI measurement of Raji B cell tumor burden over time following infusion with high-dose CAR-T cells expressing the indicated CARs. Means ± s.d are shown. Results are pooled from 2 independent experiments. Means were compared by one-way ANOVA corrected for multiple comparisons. ****, P < 0.0001; ***, P < 0.001; *, P < 0.05; ns, not significant (P > 0.05).

We tested whether 3D CARs behaved similarly to CARs elongated using the mouse CD4 spacer by comparing triggering between 0D and 3D CAR-T cells. As with αGFP-CD4 CAR, 3D CARs were minimally phosphorylated in response to stimulation with GFP-Thy1. Activation of PLCγ1, a key mediator of T cell Ca^2+^ influx and hallmark of T cell activation, was also undetectable in 3D CAR-T cells, indicating abrogation of downstream signaling (Fig. 4B). This resulted in drastically reduced (∼7 fold) cytokine secretion (Fig. 4C), but as with CD28 CARs, anti-tumor cytotoxicity was much less affected (∼2-fold) (Fig. 4D). Indeed, minimal differences in effector responses were observed between CD28 and 4-1BB CAR-T cells (Fig. 4F,G). Conjugate formation of 0D or 3D CAR-expressing CAR-T cell and target cells, as measured by a flow-cytometry-based adhesion assay, was also unaffected by elongation (Fig. 4E).

Our *in vitro* testing of titin-derived elongated CARs demonstrated, in a second CAR system, that elongation with rigid IgSF spacers results in reliable reduction of CAR triggering and cytokine secretion, while preserving cytotoxic function. Since this functional profile, of low cytokine secretion but effective tumor cell killing, are clinically desirable CAR-T cell phenotypes, we next tested whether these *in vitro* CAR-T cell profiles were reflected in anti-tumor responses in a preclinical NSG mouse leukemia/lymphoma model. We first seeded NSG mice *i.v.* with 5 x 10^5^ Raji B cells expressing GFP-Thy1 and luciferase infused, followed 5 days later with *i.v.* infusion of 10^6^ (low dose) or 10^7^ (high dose) CAR-T cells for measuring acute serum cytokine release, or longer-term monitoring of tumor burden by IVIS, respectively (Fig. 4H).

Measurement of circulating cytokines in serum of NSG mice 24 hours after CAR-T cell infusion, confirmed that CAR T cells expressing 3D CARs produce ∼20-50-fold less IFNγ, TNFα and IL-2 than OD CAR T cells (Fig. 4 I, J, and K). Strikingly, control of tumor cell burden and survival was very substantially improved in 3D vs 0D 4-1BB CARs, suggesting that the combination of a blunted acute inflammatory cytokine release and moderated CAR-T cell activation, by CAR elongation, produced functionally superior CAR-T cells (Fig. 4 L.M).

## Discussion

Here, we show that increasing CAR-antigen complex span attenuates CAR triggering, by increasing intermembrane distance and CD45 access to antigen-engaged CARs, resulting in attenuation of CAR triggering and CAR-T cell activation. However, weak triggering by elongated CARs supported cytotoxic granule polarization and exocytosis, resulting in relatively preserved anti-tumor cytotoxicity, but not NFAT activation and cytokine production, resulting in drastically reduced inflammatory cytokine release. These results reveal a hierarchy in cell-intrinsic signaling thresholds for CAR-T cell effector responses, in which CAR-T cell cytotoxicity is triggered at orders of magnitude less CAR signaling than other T cell effector responses. These hierarchical signaling thresholds for cytotoxicity and cytokine release are T cell-intrinsic and not a result of the non-native CAR signaling domain configuration, as they are also observed in human CD8+ T cell clones, in response to cognate pMHC antigens (Valitutti et al., 1996).

In this study we have used well-understood mechanistic principles of immunoreceptor signal transduction (Choudhuri and van der Merwe, 2007) to manipulate CAR signaling to produce clinically useful outcomes. In particular, our results addresses a key problem in CAR-T cell immunotherapy – the development of acute cytokine release syndrome (CRS), that occurs in as many as 40-70% of patients receiving CAR-T cell therapy (Brudno and Kochenderfer, 2019), triggered in part by inflammatory cytokine production by activated CAR-T cells – leading to local inflammation. This results in endothelial cell and macrophage activation, which in turn release IL- 1, IL-6, IL-10 and TNFα - key mediators of CRS pathogenesis (Shimabukuro-Vornhagen et al., 2018). We show *in vitro* and in a leukemia/lymphoma mouse model, that CAR elongation can dramatically reduce inflammatory cytokine secretion, while preserving cytotoxic function, resulting in low levels of CAR-derived cytokines in the circulation.

Our results provide proof-of-principle that intermembrane distance at the immunological synapse is a tunable design element that can be engineered into existing CARs for improving CAR-T cell efficacy and safety.

## Acknowledgements

This work was supported by NIH grants K99AI093884, R00AI093884 and R01AI134999 to K.C.. We thank the University of Michigan Biomedical Research Core Facilities and Rogel Cancer Center (NCI award P30CA046592) for support with microscopy and flow-cytometry/cell-sorting resources.

## Author contributions

K.C and conceived of the study, K.C and F.L. designed the study, F.L. performed experiments, K.C. and F.L. wrote the manuscript.

## Data availability statement

All data associated with the present study are available from the corresponding authors upon request.

## Methods

### Primary cells and Cell lines

Human PBMC were purchased from New York Blood Center (NYBC) as ‘leukopaks’. Raji cell and HEK293T cell line were originally obtained from ATCC. LS174T cell line, Phoenix-AMPHO cell line and CHO-K1 cell line were purchased from ATCC. Raji cells and CHO cells were cultured in complete RPMI-1640 (Gibco, 10% FCS, 2mM GlutaMAX), and HEK293T cells, Pheonix cell and LS174T cells were cultured in complete DMEM (Gibco, 10% FCS, 2mM GlutaMAX).

### Recombinant constructs

CAR constructs were cloned into the SFG retroviral vector, that has a chimeric cytokine receptor (4αβ) upstream of CARs to selectively expand CAR-T cells using IL- 4 (Wilkie et al., 2010). Coding sequences for 4αβ and CAR were separated with a furin cleavage site and T2A peptide for stoichiometric co-expression. Camelid anti-GFP nanobody, anti-CEACAM5 BW431/26 scFv and anti-CEACAM5 MFE23 scFv were used as antigen targeting domains, followed by the human CD28 transmembrane segment, followed by, CD28 or 4-1BB signalling domains, fused to the three ITAMs domains of TCRζ. Spacer sequences were inserted between the antigen-targeting domain and the CD28 transmembrane domain. All fragments were synthesized as gBlocks (IDT) and subcloned into XhoI/NotI linearized SFG-4αβ by In-fusion cloning (Clontech). To co-express PD-1 with CARs, 4αβ was substituted for full-length PD-1 by cloning PD-1 cDNA gBlock into NcoI linearized SFG-CAR constructs.

To express membrane tethered target antigens, gBlocks or PCR products encoding the target antigens were cloned into XhoI/NotI linearized pHR-sin lentiviral vector. To express membrane tethered monomeric eGFP (A206K), human β2M signal peptide and mouse H2-D1 stalk/transmembrane/cytoplasmic tail were placed at the 5’ and 3’ end of eGFP respectively.

Spacers were inserted between eGFP and stalk region by ligating the corresponding gBlocks with the NotI linearized pHR-eGFP construct.

### Cell transduction

For retroviral transduction of human T cells, Phoenix-AMPHO (ATCC) were transfected with SFG-CAR constructs using lipofectamine 3000 (ThermoFisher), virus contained medium (VCM) was collected 24/48hrs later, mixed with 5μg/ml polybrene (Sigma) and 50u/ml IL-2, then added to human PBMC pre-stimulated for 48hrs by plated bound anti-CD3ε/anti-CD28. A spinduction at 1000g for 1hr at 32°C was done to increase the transduction efficiency. VCM was substituted with complete RPMI-1640 (Gibco, 2mM GlutaMAX and 10% FCS) with 100ng/ml hIL-4 (Biolegend) 2days post transduction. CAR expressing cells were enriched by MACS method using EasySep™ FITC Positive Selection Kit II (Stemcell) plus FITC conjugated anti-VHH (for anti-GFP CAR-T) or anti-Flag (for scFv CAR-T) antibodies, and cultured for 7-10 days with 100ng/ml hIL-4. Then CD4^+^ and CD8^+^ CAR-T cells with desired surface CAR expression levels were separated by a FACS sorter (Sony SH800). A representative process of CAR-T cell MACS and FACS was shown in Supplementary figure 1A.

For lentiviral transduction of CHO and Raji cells, HEK293T cells were transfected with Phr-SIN expression vectors or pLenti CMV V5-LUC Blast (w567-1), with pRSV, pGAG and pVSV-G using lipofectamine 3000. VCM was added to CHO or Raji cells with 5μg/ml polybrene for 48hrs, before multiple times of FACS were done to match the surface levels of transduced ligands. A representative process of matching surface levels of GFP+ CHO and GFP-Thy1+ CHO was shown in Supplementary figure 1C. pLenti CMV V5-LUC transduced Raji cells were selected using 50μg/ml blasticidin for 2 weeks and the transduction were confirmed by detecting V5-tag expression using flow-cytometry.

### Flow-cytometry and cell sorting

To label cells for analysis by flow-cytometry and cell sorting, live cells were suspended in PBS/2% FBS buffer with a density of 0.5 million cells/ml, Fc blocked for 30min, and stained with surface fluorescence antibodies on ice for 30min. To detect CD107a expression, fluorescence conjugated anti-CD107a antibody was maintained during the T cell stimulation. To detect intracellular cytokine, T cells were stimulated in the presence of 2µM monesin and fixed/permeabilized using Foxp3/Transcription Factor Staining Buffer Set (eBioscience), before being stained with fluorescence antibodies targeting intracellular cytokines on ice for 30min. The stained cells were washed and passed through 70µm cell strainer before loading to BD LSRFortessa for analysis. The sorting was performed on Sony SH800 instrument with 100µm chip at the speed of 3, following gating singlets using FSC-H/FSC-W and SSC-H/SSC-W, cells were sorted based on the fluorescence signal indicating the expression levels of the target proteins. Cells were sorted into the cold FBS, and then transferred to culture medium to rest for 7-10 days prior to use in experiments. Sorted cells were checked for matched levels prior to experiments. The flow cytometry files were analysed by Flowjo10. Antibodies used were listed in supplementary table 2.

### Cytokine assay

To detect cytokines in the serum and the supernatant of CAR-T: target cell coculture, Cytometric Bead Array using enhanced sensitivity flex sets of human IL-2/IFN- γ/TNF- α was done following manufacturer’s instruction (BD Biosciences). Alternatively, sandwich ELISA were done using 1µg/ml capture antibody (anti-human IL-2 clone MQ1-17H12, anti-human IFN-γ clone MD-1), 1µg/ml biotinylated detecting antibody (anti-human IL-2 poly5176, anti-human IFN-γ clone 4S.B3) followed by labelling with 1:1000 diluted ExtrAvidin-Peroxidase (Sigma E2886) and developed by with 1-Step™ Ultra TMB-ELISA Substrate Solution (Thermo Scientific™).

### Cytotoxicity assay

CAR-T mediated cytotoxicity was assayed using DELFIA® EuTDA Cytotoxicity Reagents following manufacturer’s protocol (PerkinElmer). Briefly, target cells were loaded with 1:2000 diluted BATDA labelling reagent at 37°C for 15mins in complete culture medium. After 6 washes, 10, 000 target cells were cultured with CAR-T cells at indicated ratios in 96 V-bottom well for 2 hours at 37°C. 20µl supernatant were mixed with 200µl europium solution for 15 minutes with 250RPM shaking, before Time-Resolved Fluorescence (TRF) was detected by Envision plate-reader (PerkinElmer). TRF of target cell alone medium was set as spontaneous release, TRF of lysis buffer treated target cell was set as maximum release. The percentage of CAR-T mediated cytotoxicity was calculated as (Experimental Release – Spontaneous Release) / (Maximum Release – Spontaneous Release) ×100.

### Adhesion Assay

Experiments were performed as previously described (Choudhuri et al., 2005). Briefly, to assay CHO/CAR-Tadhesion, CAR-T cells were loaded with 2 μM CellTracker™ Deep Red [ex/em 630⁄650] in serum-free RPMI-1640 for 30 min at 37°C, CHO cells were loaded with 0.5 μM CellTracker™ Green CMFDA [ex/em 492/517] in serum-free RPMI-1640 for 30 min at 37°C, CHO cells with high GFP expression were detected by the fluorescence of GFP. After 3 washes and 30 minute-rest at 37°C in complete media, equal numbers of CHO cells and CAR-T cells (2×10^5^) were mixed and centrifuged at 500g for 10 min at 4°C (brake off). Cells were incubated for 10 min at 37°C, gently resuspended, and immediately analysed using a flow-cytometer (LSRFortessa, BD), the percentage of ‘double-positive’ red and green fluorescent events was analysed as a measure of adhesion between CHO cells and CAR-T cells.

### Biochemistry

Iimmunoblotting was performed as previously described (Choudhuri et al., 2005). Briefly, CAR-T cells and target cells expressing the indicated ligands were rapidly brought into contact by pulse centrifugation at 500g for 5 min at 4°C. After incubation at 37 °C for 0, 2 or 30 min, cells were lysed with chilled RIPA buffer (1% Triton X-100, 140mM NaCl, 50mM Hepes, 10mM Idoacetamide, 1×MS-SAFE Protease and Phosphatase Inhibitor (Sigma), PH7.4). For samples at the 0 timepoint, cells were lysed on ice immediately following centrifugation. The reduced lysate was separated by SDS-PAGE, transferred to nitrocellulose membranes, and immunoblotted for indicated signalling and control proteins using antibodies listed in Supplementary table 2. Blots were recorded using Odyssey® Classic Infrared Imaging System and analysed using Fiji/ImageJ.

### Supported lipid bilayers

Supported planar lipid bilayers were formed by depositing liposome on glass coverslips treated with piranha in Bioptechs flow chambers as previously described (Choudhuri et al., 2014; Dustin et al., 2007). DOPC liposomes contained 12.5 mol% NTA-DGS (freshly charged with Ni^3+^) to attach the his-tagged proteins. After blocking for 30min with 4% casein containing 10mM NiSO4, his-tagged proteins were incubated with lipid bilayer for 30min before thoroughly wash. Bilayers contained 200 molecules/μm^2^ of eGFP-12his or dark eGFP- 12his, and 300 molecules/μm^2^ of ICAM-1-10his (Sinobiologicals) or 500 molecules/μm^2^ of ICAM-1-10his-AF647 (in-house conjugated), or 100 molecules/μm^2^ PD-L1-AF647 (Sinobiologicals, in-house conjugated with fluorophore) in some experiments. Molecular densities on SLBs were determined by interpolation from calibration curves obtained as described previously.

### Confocal fluorescence microscopy

Equal numbers of chilled CAR-T cells and CHO cells were mixed and resuspended in ice-cold complete RPMI, before centrifuging at 500g×5min at 4°C to loosely pellet the cells. The cells were allowed to interact by incubating at 37°C for 20min. Then the cell pellet was gently resuspended with 3% PFA/PBS for 5 mins, and transferred onto poly-L-lysine treated coverslip for an additional 15 mins. The fixed cells attached to the coverslip were then washed, permeabilised with 0.1% saponin for 5mins, quenched with 100 mM glycine for 1hr, blocked with 5% BSA for 1hr. A serial staining protocol was used to stain the indicated target proteins with primary antibodies and the corresponding fluorescence conjugated F(ab’)2 of secondary antibodies, 1 hr incubation each step. Thoroughly washing were done after each staining. The coverslip with stained samples were mounted in ProLong™ Glass Antifade Mountant with the blue DNA stain NucBlue (Hoechst 33342) (Invitrogen).

The confocal images of conjugates were acquired on a NIKON A1 plus instrument, equipped with a 60x oil objective (NA 1.4, Plan Apo λ) and GaAsP PMT detector (NIKON). With the pinhole radius set to 20.43, AF405 or Hoechst 33342 were excited by 405nm laser and detected by a 450±25nm emission filter, AF568 was excited by 561nm laser and detected by a 595±25nm emission filter, AF647 was excited by 647nm laser and detected by a 700±25nm emission filter. DIC images were acquired using slider-mounted optics in the objective adaptor. The images were analysed in Fiji/ImageJ. To quantify NFAT1 cytoplasm-to-nucleus translocation, nucleus region was defined by the presence of the Hochest, the ratio of NFAT1 MFI in the nucleus region and outside of the nucleus region was calculated as a measure of the NFAT1 translocation. The spot with highest tubulin intensity was defined as MTOC position, and its distance to synapse was used as a measure of MTOC polarization. LAMP-1 polarization index was calculated as 1-(distance between interface and the center of gravity of granule mask)/(distance between interface and the opposite side of the T cell (Basu et al., 2016).

### TIRF imaging

The preparation of SLBs on piranha-cleaned coverslips is described in *Supported lipid bilayers*. To image CAR-T cell contact interfaces on SLBs, 10^6^ T cells suspended in HBS/HSA were introduced into heated FCS2 flow chambers (Bioptechs) and allowed to interact with ligand-bearing SLBs formed on coverslip incorporated into the chamber floor. Flow-chambers were maintained at 37°C throughout using a filament heater mount. After 2 or 15min incubation in flow-chambers, cells settled on SLBs were fixed with 3% PFA/HBS for 15 minutes to stop the interaction. To detect Talin recruitment, following fixation, cells were permeabilized, quenched and blocked in chamber using the same regents and incubation time as described in *Confocal fluorescence microscopy*. A serial staining with talin specific primary antibody and fluorescence conjugated F(ab’)2 secondary antibody was performed, with 1hr incubation each step.

The TIRF images were acquired on a Nikon TIRF-STORM inverted microscope fitted with a NIKON CFI SR HP Apo TIRF 100x objective (NA 1.49) and a EMCCD camera (DU-897U, Andor). TIRF illumination field (extending ∼100 nm above the SLB/coverslip surface) was used to excite fluorophores attached to antibodies at the T cell interface with bilayers. GFP was excited by 488nm laser and detected by 525±25nm emission filter, AF568 was excited by 561nm laser and detected by 600±25nm emission filter, AF647 was excited by 633nm laser and detected by 700±40nm emission filter. The images were analysed by Fiji/ImageJ, ROI of each cell was defined by SRIC image, and the fluorescence intensity in the ROI was measured. Pearson colocalization correlation (PCC) was used to analyse the linear correlation of the intensity of GFP and ICAM-1.

### Transmission electron microscopy

Conjugates of CAR-T cells and target cells were prepared as for fluorescence microscopy. After 5min incubation at 37°C, the cell pellet was gently resuspended and fixed with a mixture of 2.5% glutaraldehyde and 2% formaldehyde in 100mM cacodylate buffer (pH 7.2), post-fixed in 1% osmium tetroxide and *en bloc* stained with 2% aqueous uranyl acetate before being embedded in 2% agarose. After dehydration and infiltration with Spurr resin, ultrathin sections (∼50-55 nm thick) were cut and double-stained with uranyl acetate and lead citrate. Images were acquired on a JEOL JEM 1400 plus TEM with an AMT 12-megapixel NanoSprint12 CMOS camera (Advanced Microscopy Techniques). Intermembrane distance was measured in regions of where apposed membranes were aligned (parallel) and orthogonal to the imaging plane indicated by the trilaminar ‘tram-line’ appearance on both membranes.

### Mouse tumor model

NSG mice (NOD.Cg-Prkdcscid Il2rgtm1Wjl/SzJ) were purchased from The Jackson Laboratory and maintained under specific pathogen-free (SPF) housing conditions at the University of Michigan Medical School animal housing facility, under an Institutional Animal Care & Use Committee (IACUC) approved protocol (PRO00010222). Mice were maintained in accordance with local, state and federal regulations. 5 mice maximum were kept in one cage under the 12 light/12 dark cycle.

For Raji B cell tumor model, 8–10-week-old NSG mice were intravenously injected with 0.5 million Raji cells expressing GFP-Thy1 and firefly luciferase, CAR-T cells (10 million for high dose/1 million for low dose) were intravenously injected to the mice on day 5. Mouse survival was monitored daily, mice reached the defined humane endpoints (severe dehydration, weight loss and inability to move etc.) will be euthanized. Tumor growth in live mouse was monitored every 5 days by bioluminescence imaging (BLI) using IVIS Spectrum In Vivo Imaging System (PerkinElmer) after 10mins intraperitoneally injection of 150mg/Kg D-luciferin. Serum was collected by submandibular bleeding 24 hours after CAR-T cell administration. Bone marrow cells were harvested from femur bones of the mice treated with low dose CAR-T cells on day 14, lysed by RBC lysis buffer for 10min, filtered through 70µm filter, and processed for flow-cytometry analysis as described in *Flow-cytometry and cell sorting*, with a difference that the cells were Fc blocked with both human Fc blocker (Human TruStain FcX, Biolegend) and mouse Fc blocker (TruStain FcX™ PLUS (anti-mouse CD16/32) Antibody, Biolegend).

### Statistical analysis

Unpaired T-test and One-way ANOVA (with correction for multiple comparisons) were performed to compare multiple sample means using GraphPad PRISM9 software. PCC was used to analysis the colocalization between GFP and ICAM-1 or GFP and PD-L1 at the contact interface of T cells with SLB, or for quantitating colocalization of GFP and CD45 at conjugate interfaces.

